# A Branching Process to Characterize the Dynamics of Stem Cell Differentiation

**DOI:** 10.1101/016295

**Authors:** David G. Míguez

## Abstract

The understanding of the regulatory processes that orchestrate stem cell maintenance is a cornerstone in developmental biology. Here, we present a mathematical model based on a branching process formalism that predicts average rates of proliferative and differentiative divisions in a given stem cell population. In the context of vertebrate spinal neurogenesis, the model predicts complex non-monotonic variations in the rates of pp, pd and dd modes of division as well as in cell cycle length, in agreement with experimental results. Moreover, the model shows that the differentiation probability follows a binomial distribution, allowing us to develop equations to predict the rates of each mode of division. A phenomenological simulation of the developing spinal cord informed with the average cell cycle length and division rates predicted by the mathematical model reproduces the correct dynamics of proliferation and differentiation in terms of average numbers of progenitors and differentiated cells. Overall, the present mathematical framework represents a powerful tool to unveil the changes in the rate and mode of division of a given stem cell pool by simply quantifying numbers of cells at different times.

---

Developmental processes are tightly orchestrated in both space and time to ensure proper final form and function of organs and tissues. In the developing vertebrate central nervous system, a cycling progenitor cell faces three different outcomes upon division: the generation of two progenitor cells with self-renewing potential (pp division), two daughter cells committed to differentiation (dd division), or an asymmetric mode of division that produces one progenitor cell and one differentiating cell (pd division). Proliferative pp divisions dominate at early stages of development to expand the stem cell population without losing developmental potential, while later in development, dd divisions generate differentiated cells at the expenses of the progenitors pool. The asymmetric mode of division pd results in maintenance of the stem cell population, while differentiated cells are continuously produced [1, 2, 3].

The molecular mechanisms that govern the decision between each mode of division are beginning to be understood. This decision has been linked to the orientation of the mitotic spindle, the inheritance of polarity components, the distribution of cell-fate determinants during mitosis, the presence of extracellular morphogenetic signals, and the cell cycle length [4, 5, 6, 7, 8, 9, 2, 3, 10]. Here, we derive a general theoretical framework based on a branching process formalism that captures the average dynamics of proliferation and differentiation of a heterogeneous stem cell population in terms of balance between proliferative and differentiative divisions and average cell cycle duration, given the numbers of progenitors and differentiated cells at different times. The equations derived are then applied to study primary neurogenesis in the developing chick spinal cord, showing quantitative agreement with experimental data for the cell cycle length and rate of each mode of division. We also show that the rates of the three modes of division follow a probabilistic binomial distribution, allowing us to derive analytical equations for the rate of each mode of division. To further validate the model predictions, we developed a phenomenological in silico model of the dynamics of vertebrate neurogenesis, where we show that the values of average division rates and cell cycle length predicted by the theoretical model are sufficient to reproduce the dynamics of growth of the developing spinal cord obtained experimentally. Overall, our studies show that, despite the complex regulation of stem cell differentiation, the growth and differentiation dynamics of a given stem cell pool can be calculated based on simple mathematical assumptions.

## Results

### A Markov branching process to link division rate and division mode to progenitor and differentiated cell numbers

In general, a stem cell pool can be interpreted as a number of cells *P*_0_ at an initial time *t* = 0 that can be in quiescent (*P*_0_(1−γ)) or cycling state (*P*_0_γ), and where each cycling cell has a given probability [11, 12] to divide via the three potential modes of division (with rates *pp, pd* or *dd*) or to undergo apoptosis with a rate ∅ (*pp* + *pd* + *dd* + ∅ = 1). A schematic of these potential choices is shown in Fig. 1A. Under these assumptions, the system dynamics can be characterized by a time dependent supracritical Markovian branching process [13, 14], where the number of progenitors at an arbitrary time *t* can be obtained based on the following equation (detailed step-by-step derivation of the equations used in this section is shown as Supplementary Material)[15, 16]:

#### Significance

We present a theoretical framework to predict the dynamics of division rates and division times in stem cell populations based on the numbers of progenitors and differentiated cells at different times. The theory is applied of understand the overall dynamics of proliferation and differentiation during vertebrate neurogenesis. Overall, our studies show that, despite the complex regulation of stem cell differentiation, the growth and differentiation dynamics of a given stem cell pool can be calculated based on simple mathematical assumptions. The present mathematical framework constitutes a valuable tool to extract relevant data from complementary experimental approaches, and that its generality and simplicity ensures its straightforward application across multiple disciplines in the field of stem cell research. DGM: Designed research, performed research, wrote the manuscript.

Reserved for Publication Footnotes

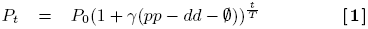

where *T* is the average cell cycle length, *γ* is the ratio of cycling cells within the population, *pp* and *dd* are the average probabilities for symmetric proliferative or differentiative division, correspondingly, while ∅ is the rate of apoptosis. The value *pp* − *dd* − ∅; can be identified as the average number of newly generated stem cells produced per division (*pp* − *dd* − ∅ = 1 corresponds to all divisions being proliferative, while *pp* − *dd* − ∅ = −1 means that all progenitor cells undergo differentiation or apoptosis). Three modes of development can be defined for the progenitor population: supra critical for *pp* − *dd* − ∅ > 0 (Fig. 1B, the number of progenitor cells increases monotonically), homeostatic for *pp* − *dd* − ∅ = 0 (Fig. 1C, where differentiated cells are produced in a linear fashion), and subcritical for *pp* − *dd* − ∅ < 0 (Fig. 1D, the number of progenitor cells reduces and the self-renewing potential of the system eventually extinguishes with all cells either differentiated or dead). Based on the same assumptions, we can derive an equation for the number of differentiated cells *D* produced at any time *t*, starting from a initial pool of differentiated cells *D*_0_:

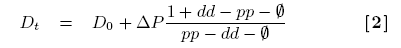

that does not depend explicitly on time *t* or the cell cycle length *T*, only on the number of progenitors generated Δ*P* = *P*_*t*_ − *P*_0_. Complementarily, the total number of cells that undergo apoptosis (Ψ) after a given time can be calculated as:

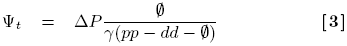

**Fig. 1.**
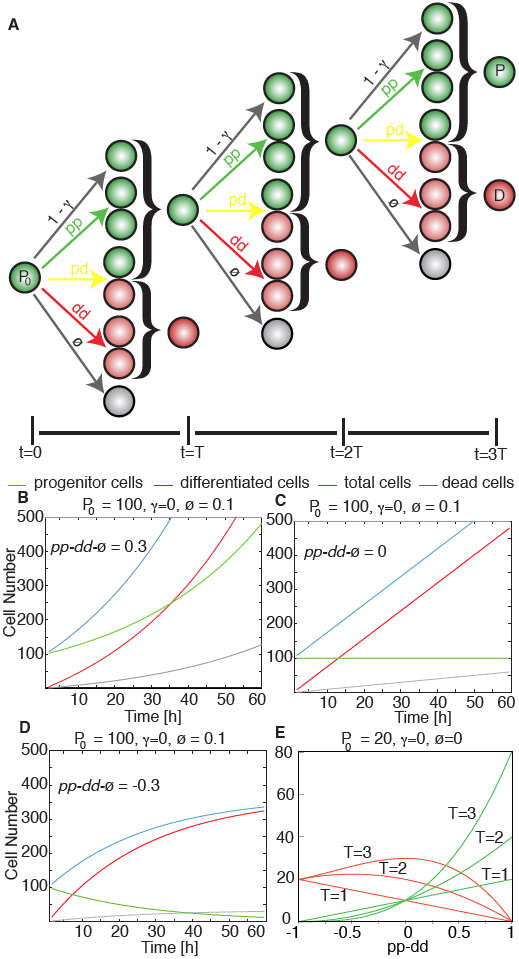
A Mathematical Model to Describe Stem Cell Populations. (A) Scheme of the branching process for stem cell behavior where a initial pool of progenitors *P*_0_ undergoes three rounds of cell division facing several potential outcomes to give a final number of progenitors *P* and differentiated cells *D*. These potential outcomes correspond with the different modes of divisions with rates *pp, pd* or *dd*, apoptosis with rate ∅, or quiescence with rate (1-*γ*). (B-D) Dynamics of progenitors and differentiated cells for different situations of growth (B, *pp* − *dd* −∅ > 0), homeostasis (C, *pp* − *dd* −∅ = 0) and reduction (D, *pp* − *dd* −∅ < 0) of the progenitors pool. (E) Solution of the model equations depending on the value of *pp* − *dd* for three time points. Dependence on ∅ and *γ* can be found as Supplementary Material.

Altogether, eqs. 1-3 allow us to directly obtain the final numbers of progenitors, differentiated and dead cells at any given time depending on the value of *pp* − *dd* − ∅ and *T*, as shown in Fig. 1E for three different multiples of the average cell cycle length *T* at fixed *γ* and apoptosis rate ∅. Supplementary Fig. 1 corresponds to plots of predicted cell numbers for varying numbers of quiescence and apoptosis rate ∅.

Eqs. 1-2 can be simply rewritten to directly obtain the proliferation rate and the cell cycle:

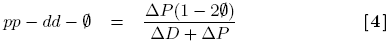

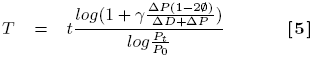

being Δ*D* = *D*_*t*_ − *D*_0_. Based on these equations, given the numbers of progenitors and postmitotic differentiated cells in a stem cell population at two time points, the value of the average rate of proliferation *pp*−*dd*−∅ and the cell cycle *T* can be calculated independently of each other, simply based on numbers of progenitors and differentiated cells at two given time points. This is of experimental relevance, since measurements of the rates of each mode of division and cell cycle are often indirect and complex to perform, while numbers of progenitor and differentiated cells can be easily quantified by immunostaining against molecular markers for each cell state. In addition, the rate of apoptosis can be determined based on active Caspase3 immunostaining, while the number of quiescent cells can also be determined experimentally using cumulative BrdU labeling [17, 2, 3]. In the next section, we apply this mathematical framework to the study of the dynamics of vertebrate neurogenesis using quantification of progenitors and differentiated cells in transversal sections of the developing chick spinal cord.

### Two waves of proliferation and differentiation while the developing spinal cord grows monotonically

The study of the dynamics and the molecular mechanisms that orchestrate the formation of the central nervous system is one of the main topics in developmental biology. In vertebrates, neural progenitor cells are organized in a pseudo stratified neuroephitelium (the neural tube) and undergo several rounds of symmetric and asymmetric divisions to generate subsets of differentiated neurons and glial cells in a process highly regulated by a number of interacting intracellular and extracellular signals and morphogens [2, 3, 18]. Vertebrate neurogenesis is also coupled in space and time with the cell cycle via interkinetic nuclear migration (INM), where DNA synthesis (S-phase) occurs at the basal side while mitosis (M-phase) occurs at the apical-most region of the neuroepithelium. When committed to differentiation, cells de-attach from the apical region and migrate basally out of the ventricular zone towards the mantle zone [19].

To study the balance between proliferation and differentiation during spinal cord development using the previous model, we proceed by quantifying the numbers of progenitor cells and differentiating neurons in transversal sections of chick embryos at different stages of development. To do so, sections are stained with antibodies against molecular markers of progenitor cells (anti-Sox2) and differentiating neurons (anti-HuC/D). Fig. 2A presents representative images of spinal cord sections obtained by confocal microscopy at various developmental stages (expressed in hours post-fertilization; HPF).

**Fig. 2.**
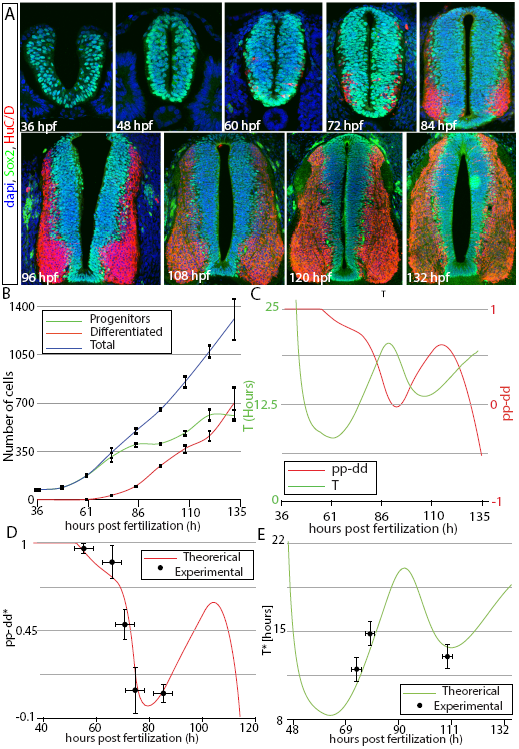
Dynamics of developing spinal cord shows two waves of proliferation. (A) Confocal snapshots of developing spinal cord sections at different HPF showing staining for progenitors (green) and differentiated (red) cells. (B) Quantification of the number of progenitor and differentiated cells overtime in one hemisection of the developing spinal cord (both left and right sides of developing spinal cord are symmetric in terms of cell numbers). Lines correspond to cubic interpolation between experimental points as a guide to the eye. Measurements were performed in at least 10 spinal cord sections from at least 3 different embryos for each time point. Error bars correspond to the standard error of the mean value for multiple independent repeats of the experiment. (C) *pp* − *dd* (red) and *T* (green) values extracted from the model equations suggesting correlation between cell cycle length and rate of neurogenesis. (D) Comparison of experimental data (dots) and theoretical (line) predictions for the rate of differentiation *pp* − *dd*\* after time correction (see Materials and Methods and Supplementary Fig. 2a). (E) Comparison of experimental data (dots) and theoretical (line) predictions for the cell cycle *T*\* corrected to take into account cells leaving teh section (see Materials and Methods and sup. Fig. 2B). Error bars in (E-D) correspond to the standard error of the mean. Experimental data obtained from [3, 26, 27].

**Fig. 3.**
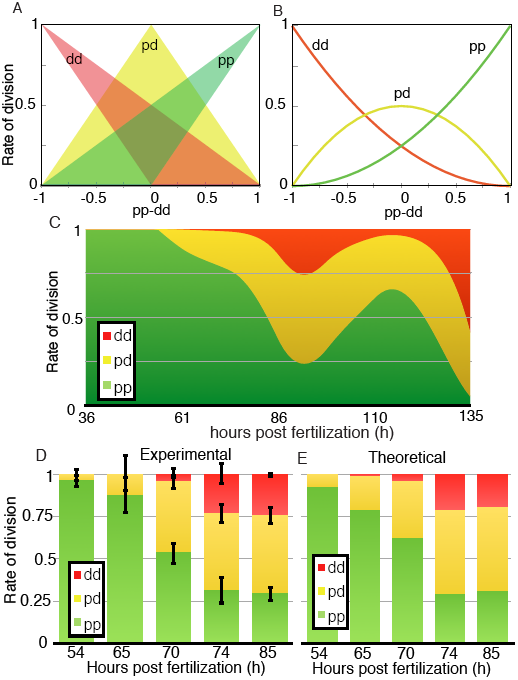
Experimental data for the division rates correlates with the binomial distribution hypothesis. (A) Possible ranges of values for each mode of division *pp*, *pd* and *dd* for each value of *pp* − *dd*. (B) Exact solution of the model equations for each mode of division when considering the binomial approach for the generation of each division mode. (C) Model predictions for each mode of division based on the experimental quantification. (D) Experimental [3] and (E) theoretical values for each rate of division at the same times given by the binomial distribution hypothesis.

Average values for progenitor and differentiated cell numbers from multiple independent repeats of the experiments are shown in Fig. 2B. At the time of dorsal spinal cord closure around 36 HPF, all cells are Sox2+ progenitors (Figs. 2A,B, green). Between 36 and 49 HPF, the number of neural progenitors (and thus, of total cells) remains constant, what is most likely due to the delamination of neural crest cells from the dorsal part of the developing spinal cord during the first stages of development [20]. From 49 HPF, the progenitor population expands until 86 HPF, followed by a regime where the number of progenitors remains constant, until 96 HPF. Then, another regime where the population of progenitors increases at 108 to 120 HPF, followed by a final regime of progenitor population homeostasis from 120 to 132 HPF. Unlike the number of total cells (Fig. 2B, blue line) which grows monotonically from 49 HPF onwards, the number of differentiated cells (Fig. 2B, red line) increases in two phases. Neuron production starts around 60 HPF, with the first differentiated cells being observed in the ventral region of the spinal cord section, reflecting the early generation of motor neurons [2, 21, 18]. From 72 HPF, differentiating cells appear along the whole dorsal-ventral axis, evidencing the generation of the different classes of interneurons from ventral and dorsal progenitor domains [22, 3, 23]. This first phase of high rate of neuron production slows down around 110 HPF, while another phase of increased differentiation begins around 120 HPF. This second wave of differentiation may correspond to the generation of late classes of dorsal interneurons (dIL) [24, 25]. Although the dynamics of the balance between growth and differentiation are not homogeneous along the dorsal-ventral axis and depends on the specific progenitor domains [2, 22, 3, 18], the average values extracted using eqs. 4-5 are suffcient to quantitatively reproduce the dynamics of growth and differentiation of the whole spinal cord section, as shown in the next sections.

### Model Predicts Nonmonotonic Changes in the Mode of Division and Cell Cycle Length

Values from total progenitors and differentiated cells are used to inform the Eqs. 4-5 to obtain the average values for *pp* − *dd* − ∅ and T at different developmental stages. To do that, we first estimate the dynamics of numbers of progenitors and differentiated cells between experimental time points using data extrapolation, as explained in the Materials and Methods section. The average rate of apoptosis has been estimated by Caspase3 immunostaining as less than 1% of the cells in each section, in agreement with recently published data [3, 18], therefore, the rate of apoptosis ∅ is assumed as negligible. Regarding the percentage of quiescent cells, experimental data show that initially all cells proliferate (*γ* = 1) to then reach a value of *γ* = 0:8 at around 100 HPF [3]. As a first approximation, change between these two data points is assumed as linear.

Values for *pp* − *dd* and *T* obtained are plotted in Fig. 2C. The model predicts an initial regime where all divisions are proliferative (i.e., *pp* − *dd* ≈ 1), followed by a regime where the number of neurogenic divisions increases at the expenses of proliferative divisions, corresponding to the generation of the first motor neurons [2]. After that, there is sharp decrease in the rate of proliferative divisions at around 70 HPF, which coincides with the developmental switch in the mode of division of the motor neuron progenitor cells reported in [2]. A second maximum in the rate of proliferative divisions occurs at 116 HPF, which then decreases again with a second wave of increased rate of differentiation taking place from 120 until 132 HPF.

Regarding the cell cycle, the model predicts an initial decrease after closure of the NT and delamitation of the neural crest cells, reaching a minimum at t=60 HPF that coincides with the initiation of differentiation. After that, the cell cycle increases until it reaches a maximum value of around 20 hours, slightly preceding the point of maximum rate of differentiation at around 90 HPF. Later on, the cell cycle shortens again at the same time as the rate of proliferative divisions increase, and reaches a second minimum at t=110 HPF that slightly precedes the second maximum in the rate of proliferation. These data suggest that increases in neurogenic divisions correlate with increases in the cell cycle length (comparison with experimental data for the correlation between mode of division and cell cycle length is addressed in the Discussion Section).

### Theoretical Predictions Correlate with Experimental Measurements of Mode and Rate of Division

Experimental quantification of the three different division rates at different developmental times has been recently achieved in the chick developing spinal cord by in ovo electroporation of single cell reporters for the mode of division [2, 3]. These markers express fluorescent proteins driven by promoters active during proliferation or differentiation, in such a way that progenitor-generating (pp and pd) divisions are identified by activation of the Sox2p enhancer driving expression of EGFP, while neurogenic (pd and dd) divisions are identified by the activation of the Tis21 promoter driving expression of RFP. Quantification of the average mode of division of mitotic cells in whole chick spinal cord hemisections has been recently performed [3]. Since our model is informed with numbers of all cells in all stages of the cell cycle (not only in M-phase), comparison with the experimental data should be performed after rescaling of the time variable t to account for the average time of the cells in the population since its generation (See Materials and Methods and Supplementary Fig. 2A for a detailed explanation). Curve in Fig. 2D corresponds to this rescaled value of the theoretical *pp*−*dd*\*, showing quantitative agreement with all available experimental values for *pp*−*dd* (dots in Fig. 2D) computed using data from [3].

Experimental quantification of the average cell cycle length in developing chick spinal cord sections can be estimated by Brdu-Edu cumulative curves [2, 26, 27, 3]. To compare this value with our model predictions, we have to take into account that the spinal cord is also growing in the anteroposterior direction, and therefore, a number of cells generated in a transversal section are contributing to the axial growth of the embryo. This number can be estimated from recent experimental data [18], that reports a 15% growth of the chick spinal cord in the anteroposterior axis after 15 hours, (see Supplementary Text and Supplementary Fig. 2B). In addition, eqs. 4-5 can be generalized to take into account this rate, showing that the mode of division does not depend on the anteroposterior growth and can be estimated from quantification of cells in fixed sections. On the other hand, the cell cycle length needs to be recalculated based on the total number of cells produced (see Supplementary Text and Supplementary Fig 2D). Comparison of theoretical predictions of the corrected cell cycle length *T*\* with previously published experimental data [26, 27, 3] is shown in Fig 2E for the three available time points.

### The rates for each mode of division fit with a scenario of independent probability of differentiation of daughter cells

As a consequence of the mathematical equivalence between *pd, pp* and *dd* (see Supplementary Fig. 4), multiple combinations of *pp, pd* and *dd* are possible for a given value of *pp* − *dd*. Fig. 3A illustrates the potential values for each mode of division depending on the value of *pp* − *dd*. In principle, the solution *pd* = 0 is possible for any value of *pp* − *dd*, while at intermediate values of *pp* − *dd, pd* can take any value between 0 and 1. Although the outcome of asymmetric or symmetric divisions may be equivalent in terms of cell numbers, the biological regulation required in both scenarios can be highly different, in terms of integrating extracellular signals to actively regulate the localization and distribution of cell fate determinants during mitosis. Due to this redundancy, in a given developmental scenario, asymmetric divisions can be favored versus purely symmetric divisions or vice versa depending on the specifics of each biological system. For instance, neurons and glial cells in Drosophila are generated specifically via asymmetric cell divisions [28], while motor neuron and interneurons generation in vertebrates is achieved using a combination of symmetric neurogenic and asymmetric divisions [2, 3].

**Fig. 4.**
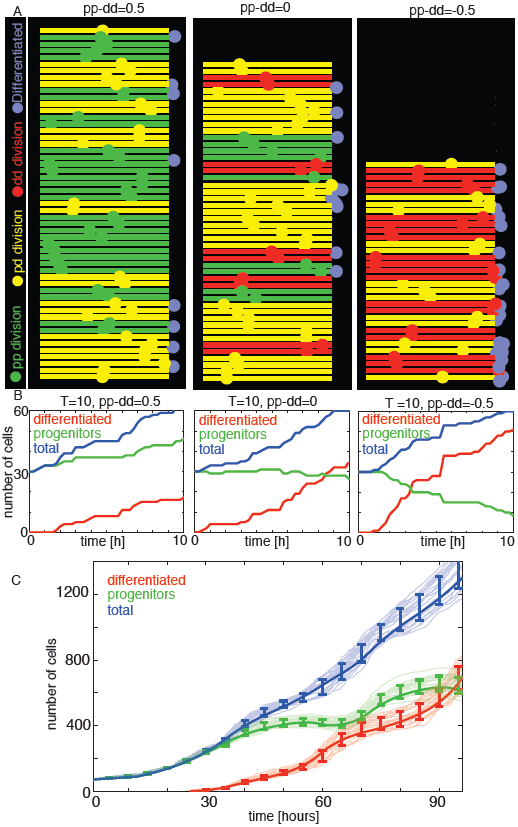
Phenomenological simulation of developing spinal cord growth reproduces the dynamics of the experimental system. (A-C) Snapshots of three simulations with constant values for the rates of division and cell cycle length, after an initial value *P*_0_ = 30, *D*_0_ = 0 showing growth (*A*, *pp* − *dd* > 0), maintenance (B, *pp* − *dd* = 0) and decrease (C, *pp* − *dd* = 0) of the progenitor population. (D-F) Dynamics of the two cell populations in the three different regimes explored. (G) Dynamics of the two populations using the values of the cell cycle length and mode of division predicted by the theoretical model. Semi-transparent lines correspond to different runs of the model. Error-bars correspond to the standard deviation of the mean over 20 different simulations. Solid lines correspond to the experimental data. Start of the simulation (0 hours) correspond to 36 HPF in the experiment.

Alternatively, if the decision between differentiation or proliferation of a given progenitor cell is purely stochastic [11, 16, 29], and the probability of differentiation of two sister cells is considered independent of each other, the probability of proliferation and differentiation of a *n* number of newborn cells can be defined as a binomial distribution of the form:

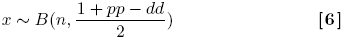

where the outcome variable *x* depends on the probability of becoming a progenitor cell, which can be written in terms of the rates of proliferative and differentiative divisions as 0 < (1+*pp*−*dd*)/2 < 1 (see Supplementary Text for a detailed explanation).

In this scenario, an asymmetric division can be identified as two independent probabilistic events (*n* = 2) that result in one progenitor and one differentiated cell (*x* = 1). This way, we can derive the probability of *pd* division as the probability of one of the daughter cells to maintain the progenitor fate while the other acquires a differentiated fate as:

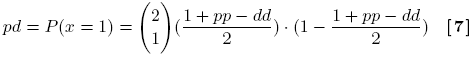

that, taking into account that *pp* + *pd* + *dd* = 1, it allows us to obtain expressions for the rate of each mode of division based on the value *pp* − *dd* (see Supplementary Text):

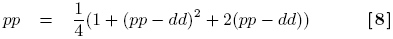

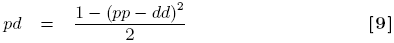

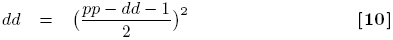

These values are plotted for each value of *pp* − *dd* in Fig. 3B. This way, if a given experimental system exhibits a rate of pd divisions higher that the value predicted by eq. 9, we can assume that the probabilities of differentiation of sister cells are not independent, in such a way that the system actively regulates the distribution of cell fate determinants to favor asymmetric divisions. On the other hand, lower values of *pd* evidence a preferential use of a symmetric distribution of cell fate determinants to favor proliferative or neurogenic symmetric divisions versus asymmetric *pd* divisions.

When we consider this particular scenario to estimate the rates for the three modes of divisions overtime (Fig. 3C), we see that purely proliferative divisions initially dominate, while most of the neurogenic divisions are predicted to be mainly asymmetric. The number of symmetric neurogenic dd divisions is predicted to peak at 92 HFP, corresponding to around 25% of all divisions, and at 136 HPF, corresponding to around 50% of all divisions occurring at these stages.

Comparison of these values with the experimental data extracted from [3] at five time points (Fig. 3D experimental, to be compared with Fig. 3E theoretical) shows quantitative agreement with the calculations derived from the binomial distribution of cell fate. This evidences that the system does not favor asymmetric versus symmetric divisions, or vice-versa, and therefore, the rate of each mode of division can be calculated assuming a probabilistic scenario of both daughter cells deciding to proliferate or differentiate stochastically and independently of each other. In consequence, using eq. 4 and the relation *pp* + *pd* + *dd* = 1, exact rates for each mode of division *pp, pd* and *dd* can be calculated analytically as (see Supplementary Text):

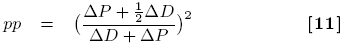

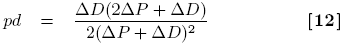

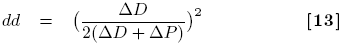

that together with eq. 5, fully characterize the dynamics of the system in terms of rates for each mode of division and cell cycle length based simply on quantification of progenitors and differentiated cells at different time points.

### Simulations of spinal cord development reproduce the experimental dynamics

To validate the theoretical prediction for the values of *pp, pd, dd* and *T*, we developed a phenomenological *in silico* model of a spinal cord hemisection, where we define cells that proliferate and differentiate with a given probability. Cells are confined in a simulated ventricular zone and move from apical to basal, mimicking the mechanism of INM [19]. This way, mitosis occurs at the apical boundary, while differentiated cells de-attach apically, migrate laterally and leave the ventricular zone. To mimic the *in vivo* cell division markers [2, 3], cells resulting from a *pp, pd* and dd division are labeled as green, yellow and red, respectively. Differentiated cells are labeled as magenta. Fig. 4A shows three different simulations with three different constant values of *pp* − *dd* and cell cycle length, where a initial set of 30 progenitor cells is allowed to develop during a single cell cycle. To mimic cell variability and the stochastic nature of cell differentiation [16, 30, 29], cell cycle length for each cell is obtained from a gamma distribution with mean equals to *T*, and standard deviation of 30% of the mean. Quantification of the three simulations is shown in Fig. 4B, where we can see the three different regimes of growth (*pp* − *dd* > 0), homeostasis (*pp* − *dd* = 0) and reduction (*pp* − *dd* < 0) of the initial population of progenitor cells. Time-lapse movies of the system for different constant differentiation probabilities are available as Supplementary Material.

When we inform the simulation with the values of *T* and *pp* − *dd* predicted by the theoretical model (Fig. 2C), the resulting dynamics of development for each cell state is shown in Fig. 4C, where light blue (total), red (differentiated) and green (progenitor cells) lines correspond to 20 independent runs of the model. The initial number of progenitors is set at 70 cells, corresponding to the experimental data in Fig. 2B at 36 HPF. Experimental values for the dynamics of each population are also shown as dark blue, red and green, to illustrate the quantitative agreement between experimental data and the simulation. This evidences that, despite cell-to-cell variability, stochasticity in the differentiation process, and heterogeneity in the dynamics between the different progenitor subdomains, the average values of *T*, *pp*, *pd* and dd predicted by eqs. 4-5 fully capture the dynamics of differentiation and proliferation during spinal cord development.

## Discussion

During neural development, the process of differentiation of progenitor cells is tightly regulated by extracellular signals and morphogens that ultimately trigger the expression of proteins specific for each cell state [31, 32, 2, 3]. Previous theoretical approaches to study stem cell dynamics focused in the general aspects of stem cell biology [15, 33], applied to colon crypts [13, 14], hematopoietic [16, 34], or cancer stem cells [35]. In the context of motor neuron generation [2], we recently developed a mathematical model based on chemical kinetics formalism that predicted a switch in the division mode in synchrony with changes in Shh activity.

Here, we take a complementary approach by developing a mathematical framework based on a branching process formalism to study the dynamics of proliferation and differentiation of a stem cell pool that provide us with analytical solutions for the average mode and rate of division of the population. The theory assumes that cells can either self-renew, differentiate, undergo apoptosis or remain in a quiescent state, and that this decision can change overtime during organ development. We assumed a linear change for the amount of quiescent progenitors P(1-*γ*) up to a 20% of the total progenitors at 100 HPF (based on data from [3]). Importantly, the value of *pp* − *dd* does not depend on *γ*, as shown in eq. 4, so this assumption does not impact the calculation for the rates of the mode of division. We also developed a generalized equation for the cell cycle length to account for the cells that contribute to the axial growth of the spinal cord (see Supplementary Text), that shows quantitative agreement with experimental data for the division rate obtained using Brdu-Edu cumulative curves [26, 27, 3].

Our results show that the average cell cycle increases with the increasing of neurogenic divisions (see Fig. 2C). This is in agreement with experimental data [36, 37, 38, 39, 40, 41, 42, 43, 44, 45], but not with other recent studies in spinal cord, retina and cortex [46, 3, 2, 47, 48] that report a reduction in the cell cycle length, due mainly to changes in the S-phase or G2 of the cell cycle [2, 26], or even no impact in cell cycle length associated to an increase in neurogenesis [26, 27, 49]. Comparison of the theoretically predicted curves for *pp* − *dd* (Fig. 2D) and *T* (Fig. 2E) developed to be compared to the experimental data (see Methods Section and Supplementary Material for a detailed explanation of the generation of these curves), is shown as Supplementary Fig. 2D. Superimposition of the curves shows regimes where the lengthening of the cell cycle correlates (yellow) or anti-correlates (blue) with increasing differentiation. Therefore, when comparing values from Brdu-Edu accumulation and differentiation of cells in mitosis, our data suggests that the correlation between changes in mode and rate of division depends strongly on the developmental stage that is being studied.

Recent experiments evidence that, in the context of the spinal cord, the regulation of the mode of division precedes alteration in the length of cell cycle [3]. On the other hand, our data (Fig 2C) shows changes in cell cycle slightly preceding changes in the mode of division. This is due to the fact that variations in the probability of differentiation of daughter cells impact the length of the cell cycle of the mother, as discussed above. This way, variations in the cell cycle length occur before daughter cells are born, and therefore, cell cycle alterations are detected prior to changes in the mode of division, when calculated based on the number of cell in the population.

The fact that the experimental data for the mode of division fits to a binomial distribution suggests an scenario where the extracellular signals sensed by the mother cell set the probability of differentiation (and not the mode of division) for the two daughter cells. This way, after a division event, the two daughter cells inherit the same probability of differentiation, but the final decision is stochastic and independent of each other. This scenario only applies to systems where the two daughter cells inherit the same probability of differentiation, i.e, cell fate determinants are evenly distributed during mitosis. This way, when wild type mitotic spindle orientation is disrupted via up-regulation or down-regulation of regulatory proteins, the amount of cell fate determinants inherited by both daughter cells is different, and the ratios between the three modes of division should not correspond to the binomial approximation. This is consistent with recent experiments in the chick spinal cord, where reduction of Inscuteable results in an overall increase of mitosis with oblique spindle orientation and increased neurogenesis [4], while randomization of the orientation of the mitotic spindle do not impact the average rates of differentiation [8].

The quantitative agreement between the *in silico* phenomenological simulation and the experimental data is presented as an additional validation of the predictions obtained by the analytical equations, showing that despite the inherent cell-to-cell variability and the different timing of differentiation of progenitor subdomains [2, 3, 18], the dynamics of growth and differentiation of the developing spinal cord is well captured by the average values of cell cycle and mode of division predicted by eqs. 4-5.

In conclusion, we believe that the present theoretical framework can be used not only to determine the values of cell cycle length and different division rates in wild type conditions, as illustrated in the present manuscript as a proof of concept. A potential more relevant use of the model will be to quantitatively characterize the influence on cell cycle length or the rate of a particular mode of division in a given developmental system, after up-regulation or down-regulation of a protein of interest, compared to the control situation. In conclusion, we believe that the present mathematical framework constitutes a valuable tool to extract relevant data from complementary experimental approaches, and that its generality and simplicity ensures its straightforward application across multiple disciplines in the field of stem cell research.

## Materials and Methods

*. Model equations Models equations have been solved numerically combining programmed datasheets in Numbers© for Mac and in-house developed Matlab© scripts (The Mathworks© Natick, MA). Code available as Supplementary Material.

### Sample preparation, Immunohistochemistry and Image acquisition

White-Leghorn chick embryos were incubated at 38 *C*^0^ and then fixed at the required HPF for 1-2 hours at RT in 4% paraformaldehyde in PBS. Interlimb sections where dissected and embedded in 5% agarose to be sectioned in a Leica ©vibratome (VT 1000S). Immunostainings were performed following standard procedures using the following monoclonal antibodies: Sox2 (Invitrogen© A-48-1400); HuCD (Molecular Probes© A-21271). Single-and double-label analyzes were performed using Alexa488-, Alexa555 (Molecular Probes©) and Cy5-conjugated (Jackson Immuno Research Inc.©) secondary antibodies. Nuclear staining performed using Dapi, Sigma© D9542. Images were acquired in a Leica SP5 confocal microscope. The sections for quantification result from superimposing three confocal images taken 5 *μm* apart in the head-to-tail axis, so the numbers of cells quantified in each section correspond to a section of 10 *μm*.

### Quantification of cells

Quantification of the number of cells in each state has been performed using ImageJ scripts in the following way: first we identified each nuclei using Dapi staining. Next, we quantify the number of nuclei corresponding to differentiated cells suing the HuC/D cytoplasm of postmitotic differentiated cells. Then we subtract the number of differentiated cells from the total cells to obtain the number of progenitor cells. Cells stained with both HuC/D and Sox2 are considered as differentiated. Mean and standard deviation are obtained from at least 3 different embryos and at least 10 different inter limb sections for each data point.

### Data extrapolation and curve fitting

Values for the numbers of cells between experimental data points were estimated using extrapolation with Matlab© algorithms (The Mathworks©, Natick, MA). To obtain the curves of Fig. 2C-D, we used a corrected “spline” interpolation to generate smooth-changing curves. A more sharp fitting (cubic interpolation) was used to obtain the values of *pp* − *dd* and *T* to inform the simulations of the neural tube dynamics of Fig. 4C. The “spline” interpolation values generated data of numbers of progenitors cells and proliferating cells with an error of around 3%, while the “cubic” interpolation error was around 0.1% compared to the experimental data.

### Generation of theoretical curves to be compared with experimental data

Experimental values for the mode of division [3] corresponds to data for mitotic cells, while the theoretical predictions are based on total numbers of cells (at any given point in its cycle), i.e., each cell in the population is monitored a certain time after the division event that that generate it. To correlate experimental and theoretical data, the time variable is rescaled to subtract the time for each cell that has passed since its generation, which corresponds to a value of is *T*/2 when we average over an asynchronous population of cells, (see Supplementary Fig. 2A). This value can be identified as the average time that has passed after the generation of all the cells in the population.

To compare experimental and theoretical values of the cell cycle length, we have to take into account that a number of newborn cells in a section contribute to the anteroposterior growth of the embryo [18] (see Supplementary Text). We developed generalized equations that account for this rate of cells that contribute to the axial growth of the spinal cord. The values of progenitors and differentiated cells obtained based on the data of anteroposterior growth by [18] are plotted in Supplementary Fig. 2C. We also take into account that the BrdU cumulative curves are calculated by averaging the value of the cell cycle during the length of the experiment (3 hours until reaching plateau in Ref. [26], and 8 hours in Refs. [27, 3], therefore the value measured corresponds to an average of the cell cycle length during the length of the accumulation experiment. Therefore, the experimental data has to be compared to a theoretical curve where the same average is performed (each point corresponds to the average value of the 6 previous hours). This data is presented in Fig. 2E.

### Simulations of developing spinal cord section dynamics

Simulations were performed using an in-house developed Matlab script. Code available as Supplementary Material. Cells are identified as numerical entities organized in a vertical axis that travel a fixed distance “d” from apical and basal regions with a speed “V” determined by the cell cycle length “T” (V=d/T), in a process that mimics interkinetic nuclear migration INM. Cell decisions between proliferation and differentiation are based on a stochastic algorithm with probability defined by the values pp, pd and dd obtained by the theoretical model (eqs. 11-13). Cell cycle length (i.e., the speed of each nuclei traveling apical to basal) is defined as a random value within a gamma distribution with mean the predicted cell cycle value by the theoretical model (T) using eq. 5 and standard deviation of 30% of the mean value.

Cells arriving to the apical region (left side of the simulation) undergo mitosis. Each newborn cell is located at the vicinity of the mitosis, with a fate determined by the mode of division that took place. Cells are color coded depending on the mode of division that generated them, mimicking the colors obtained experimentally by combination of the two cell fate markers explained in the text and developed in [2]. Cells arriving to the basal domain (right side of the simulation) will behave depending on their fate: progenitor cells will revert their direction and travel back to the apical domain, while cells with a differentiated fate will leave the ventricular zone and be located at the vicinity of its exit point, labeled as purple to mimic HuC/D immunostaining. Cells are initially all progenitors uncorrelated in location and cell cycle phase. Movies of the simulation process for different parameters can be found as Supplementary Material, for *pp* − *dd* = 0.5 and *pp* − *dd* = 0.

## ACKNOWLEDGMENTS

This work has been supported by the Ministry of Science and Technology of Spain via a Ramon y Cajal Fellowship (Ref. RYC-2010-07450), a grant from Plan Nacional framework (Ref. BFU2011-30303), and a Marie Curie International Reintegration Grant from the EU (Ref. 248346-NMSSBLS), as well as financial support from the CSIC-SPAIN (JAE-DOC fellowship). We are extremely grateful to Elisa Martϭ, Murielle Saade, Gwen le Dreau, ρngeles Rabρdan and Susana Usieto for invaluable help in the preparation of the manuscript, guidance on the experimental work and technical assistance.

